# Copper nanoparticle application enhances plant growth and grain yield in maize under drought stress conditions

**DOI:** 10.1101/2020.02.24.963132

**Authors:** Dong Van Nguyen, Huong Mai Nguyen, Nga Thanh Le, Kien Huu Nguyen, Huong Mai Le, Anh Trung Nguyen, Ngan Thi Thu Dinh, Son Anh Hoang, Chien Van Ha

**Author notes:** Equally contributed (Dong Van Nguyen, Huong Mai Nguyen, Nga Thanh Le). Corresponding author: Email address (D. V. Nguyen); (C. V. Ha).

## Abstract

Abiotic stresses, including drought, detrimentally affect the growth and productivity of many economically important crop plants, leading to significant yield losses, which can result in food shortages and threaten the sustainability of agriculture. Balancing between plant growth and stress responses is one of the most important characters for agricultural application to maximize plant production. In this study, we initially report that copper nanoparticle priming positively regulates drought stress responses in maize. The copper nanoparticle priming plants displayed enhanced drought tolerance indicated by their higher leaf water content and plant biomass under drought as compared with water-treated plants. Moreover, our data showed that the treatment of copper nanoparticle on plants increased anthocyanin, chlorophyll and carotenoid contents compared to water-treated plants under drought stress conditions. Additionally, histochemical analyses with nitro blue tetrazolium and 3,3’-diaminobenzidine revealed that reactive oxygen species accumulation of priming plants was decreased as a result of enhancement of reactive oxygen species scavenging enzyme activities under drought. Furthermore, our comparative yield analysis data indicated applying copper nanoparticle to plant increased total seed number and grain yield under drought stress conditions. Our data provided the evidences that copper nanoparticle regulates plant protective mechanisms associated with drought tolerance, which is a promising approach for the production of drought tolerant crop plants.

## INTRODUCTION

Global climate changes are resulting in an increasing of extreme environmental stresses, such as drought, heat, cold, salt, flooding, which could lead to significant yield losses of agricultural production (Stevanovic et al. 2016; Wheeler and von Braun 2013). In addition to that, the world population is increasing rapidly, setting a food security for sustainable human life (Myers et al. 2017; Wheeler and von Braun 2013). Maize (*Zea mays*), also known as corn, is one of the most important crops world-wide to provide not only food and feed but bio-fuel resources. Maize was originated from Mexico about 8700 years ago (Piperno et al. 2009). The United States is largest maize producer that supplies more than 30% of global maize production (https://www.statista.com). However, maize production is also detrimentally affected by environmental stresses, including drought (Webber et al. 2018). Therefore, development of drought-tolerant maize varieties as well as methodologies are considered as the most economical and effective for the sustainable maize production.

Recent studies have revealed that the chemical compounds such as hormones, reactive oxygen-nitrogen-sulfur species, amino acids, ascorbic acid, polyethylene glycol solution and organic compounds could help plants to improve abiotic stress tolerances (Ha et al. 2014b; Hossain et al. 2015; Nguyen et al. 2017; Nguyen et al. 2018a; Salemi 2019; Savvides et al. 2016; Utsumi et al. 2019). Various chemicals have potential applications for agricultural biotechnology, and chemical application, including nanotechnology application, would be one of the most promising methods for enhancing the abiotic stress tolerance in the fields. In recent decades, nanobiotechnology is prominent chemical application. Nanobiotechnology is an integration of physical sciences, molecular engineering, biology, chemistry and biotechnology, which holds considerable promise of advances in pharmaceuticals and healthcare (Singh et al. 2017b; Thangavelu et al. 2018). In plants, recent studies reported that nanoparticles involved in various physiological and biochemical processes controlling plant growth and development as well as plant environmental stress responses (Arora et al. 2012; Regier et al. 2015). Application of nanoparticles showed effect on plant growth, yield and crop product quality (Burke et al. 2015; Ngo et al. 2014), suggesting that nanotechnology is being a promising application approach for sustainable agriculture.

Copper (Cu) is known as an essential component, which functions in regulating plant growth and development, including chlorophyll formation and seed production (Viera et al. 2019; Xue et al. 2014). Cu is one of the micronutrients needed in very small quantities by plants (Printz et al. 2016). Several important crops, including maize, are sensitive to Cu deficiency (Karamanos et al. 2004). Cu also involves in several enzyme systems that regulate the rate of many biochemical reactions in plant (Drazkiewicz et al. 2004; Singh et al. 2017a). It is also required in the process of photosynthesis, which is essential for plant respiration and assists in plant metabolism of carbohydrates and proteins (Ambrosini et al. 2018; Casimiro and Arrabaça 2015; Din et al. 2017; Regier et al. 2015; Singh et al. 2017a). Application of free metal Cu nanoparticles (nano-Cu°) showed effect on seed yield and quality in soybean (Ngo et al. 2014). However, the function of nano-Cu° in maize under drought condition is still remain unknown.

In the present study, we evaluate the role of the nano-Cu° in controlling maize growth and development as well as drought stress responses. We show that nano-Cu° priming maintains leaf water status, and chlorophyll and carotenoid content, which could contribute to enhance plant growth and biomass in maize under drought and recovery. We also reveal that nano-Cu° priming increases the superoxide dismutase (SOD) and ascorbate peroxidase (APX) enzyme activities, and anthocyanin content under drought stress conditions, which could result in reduced the excessive of reactive oxygen species (ROS) productions and therefore increased the adaptation to drought stress conditions in maize. These findings could be useful for agricultural application by the use of nano-Cu° to enhance plant biomass as well as productivity in maize.

## MATERIALS AND METHODS

### Plant growth and treatments

#### For Cu nanoparticle concentration screening assay

seeds of the Vietnamese maize elite (namely LVN10, provided by Maize Research Institute, Vietnam) were sterilized and treated with 52 (3.333 mg/L), 69.4 (4.444 mg/L) and 86.8 µM (5.556 mg/L) nano-Cu° (free metal nano-Cu° were provided by Institute of Materials Science, Vietnam Academy of Science and Technology, Vietnam, has 30 to 40 nm in size) or water (control) for 8 hours. 10 treated seeds were sown on 6-litter pot containing super mix soil. Maize plants were grown under normal conditions (800 µmol m^-2^ s^-1^ photon flux intensity, photoperiod of 12 h light/12 h dark cycle, temperature of 32°C during the light period and 24°C during the dark period, 60 % relative room humidity), and measured the plant height at 7, 14 and 21 days after sowing. Ten biological replicates were performed. After that, the aerial portion was cut and dried for 72 h at 65°C, and then measured the shoot dried weight (DW) at 7, 14 and 21 days after sowing. Five biological replicates were collected for each experiment.

#### For drought tolerant assay

6 seeds were sterilized and sown on 6-litter pot containing super mix soil for 12 days under normal conditions (800 µmol m^-2^ s^-1^ photon flux intensity, photoperiod of 12 h light/12 h dark cycle, temperature of 32°C during the light period and 24°C during the dark period, 60 % relative room humidity). Twelve-day-old plants were treated with 450 mL each pot of 69.4 µM nano-Cu° or water (control) for 2 days, and exposed to drought treatment for 7, 14 and 21 days, then collected samples for measuring plant biomass (shoot DW). Five biological replicates were collected for each experiment.

#### For plant recovery assay

Maize plants were treated and exposed to drought stress for 21 days as described above, then re-watered and continuously grown under normal condition (800 µmol m^-2^ s^-1^ photon flux intensity, photoperiod of 12 h light/12 h dark cycle, temperature of 32°C during the light period and 24°C during the dark period, 60 % relative room humidity) for 7 days and measured shoot DW. Five biological replicates were collected for each experiment.

### Grain yield assay in maize under drought

Six maize seeds were sterilized and sown on 6-litter pot containing super mix soil for 12 days under normal conditions (800 µmol m^-2^ s^-1^ photon flux intensity, photoperiod of 12 h light/12 h dark cycle, temperature of 32°C during the light period and 24°C during the dark period, 60 % relative room humidity). Plants were treated with 450 mL of 69.4 µM nano-Cu° or water control for 2 days, and exposed to drought for 21 days. Then plants were re-watered and removed plants each pot. Two plants were continuously grown under normal condition to harvest seeds. Collected ears were dried for 72 h at 65°C and then measured the seed number and dry weight. Six biological replicates were performed for each experiment.

### Determination of anthocyanin, chlorophyll and carotenoid contents, and relative water contents

Plants were grown and treated with 450 mL of 69.4 µM nano-Cu° or water control as described in drought tolerant assay. At 7, 14 and 21 days after drought stress, the fourth leaf of stressed and non-stressed maize plants were separately collected for measuring anthocyanin as previously described (Nguyen et al. 2016), and chlorophyll and carotenoid contents as described before (Mostofa et al. 2015). An amount of 0.5 g leaf sample of maize plants under control or nano-Cu° treatment conditions were extracted by shaking (200 rpm) in dark in 20 mL of extraction solution (acetone 80%) overnight at room temperature. After the extraction, 1 mL of extraction solution was used for measuring the absorbance at 470, 645 and 663 nm using the spectrophotometry (Thermo Scientific Gensys 20, USA). Chlorophyll (*a* and *b* and total) and carotenoid contents were calculated as described in (Mostofa et al. 2015). The relative leaf water content (RWC) of the fifth leaf of maize plants during drought were measured according to previously described methods (Ha et al. 2013b). Five biological replicates were measured for each experiment.

### ROS histochemical staining detection of maize leaf

Plants were grown and treated as described in drought tolerant assay. To determine the ROS accumulation during drought treatments, the fifth leaf was collected for staining assays. Samples were stained with nitro blue tetrazolium (NBT) to detect superoxide and 3,3’-diaminobenzidine (DAB) to detect hydrogen peroxide as previous described methods (Mostofa et al. 2015). Three biological replicates each experiment was performed.

### Protein extraction and enzyme activity assay

Twelve-day-old plants were treated with 450 mL of 69.4 µM nano-Cu° or water control for 2 days. Plants were then exposed to drought treatment for 7 days and collected samples for protein extraction. 0.2 g of the fifth leaf was collected and extracted total soluble protein as previously described (Bradford 1976). The bovine serum albumin was used as a standard reference. Five biological replicates were performed. For enzyme activity assays: the APX and SOD activity was measured as described in (Mostofa et al. 2015). Three biological replicates each experiment was performed.

### Statistical analysis

Data were analyzed by using analysis of variance (ANOVA) or Student’s *t*-test. Different superscripted letters within the column indicate statistically significant differences among the treatments examined according to Duncan’s multiple range test (*P* < 0.05) (using IBM SPSS software package 21.0), and asterisks indicate significant differences as determined by Student’s *t*-test (\**P* < 0.05, \*\**P* < 0.01, \*\*\**P* < 0.001).

## RESULTS

### Improvement of plant growth and drought response by nano-Cu° treatment in maize

To elucidate the function of nano-Cu° in plant growth and drought stress response, the maize plants were grown and treated with nano-Cu° under drought or normal conditions. The effect of copper nanoparticles on the growth parameters of maize under normal and drought stress conditions were investigated in terms of shoot height and biomass (shoot DW).

First, we tested the effect of nano-Cu° on maize growth using seed priming method under normal growth conditions. As showed in Supplementary Figure 1, the shoot height of 14 days after sowing and shoot biomass of 14 and 21 days after sowing of nano-Cu° priming plants (69.4 µM nano-Cu° concentration) showed greater than that of water-treated plants. The data indicate that nano-Cu° promotes maize growth, at least at 69.4 µM concentration. There are no significant differences between other tested nano-Cu° concentrations as compared with control plants (Supplementary Figure 1). Based on that data, we designed to use 69.4 µM of nano-Cu° for all further analyzed experiments.

Second, we conducted a drought tolerant assay to investigate the role of nano-Cu° on maize drought responses. Under non-stressed conditions, we found that the shoot biomass was increased by nano-Cu°-treated plants as compared to water-treated plants at 7 and 14 days after treatment (equal to 21 and 28 days after sowing) (Figure 1A-C). Under drought stressed conditions, there were significant differences of shoot biomass between nano-Cu°-treated and water-treated groups (Figure 1A-C). The nano-Cu°-treated plants showed less wilting and rolling leaves as compared with water-treated plants after 7 days of drought treatment (Figure 1B). The nano-Cu°-treated plants exhibited less effect by drought indicated by greater shoot biomass in comparison with water-treated plants (Figure 1C). As compared with non-stressed plants, shoot biomass showed reduced 78 and 87% of nano-Cu°-treated and water-treated plants, respectively (Figure 1D). These results indicated that nano-Cu° priming in maize reduced negative impact of drought, which may result in recusing plant productivity under drought stress conditions.

**Figure 1.**
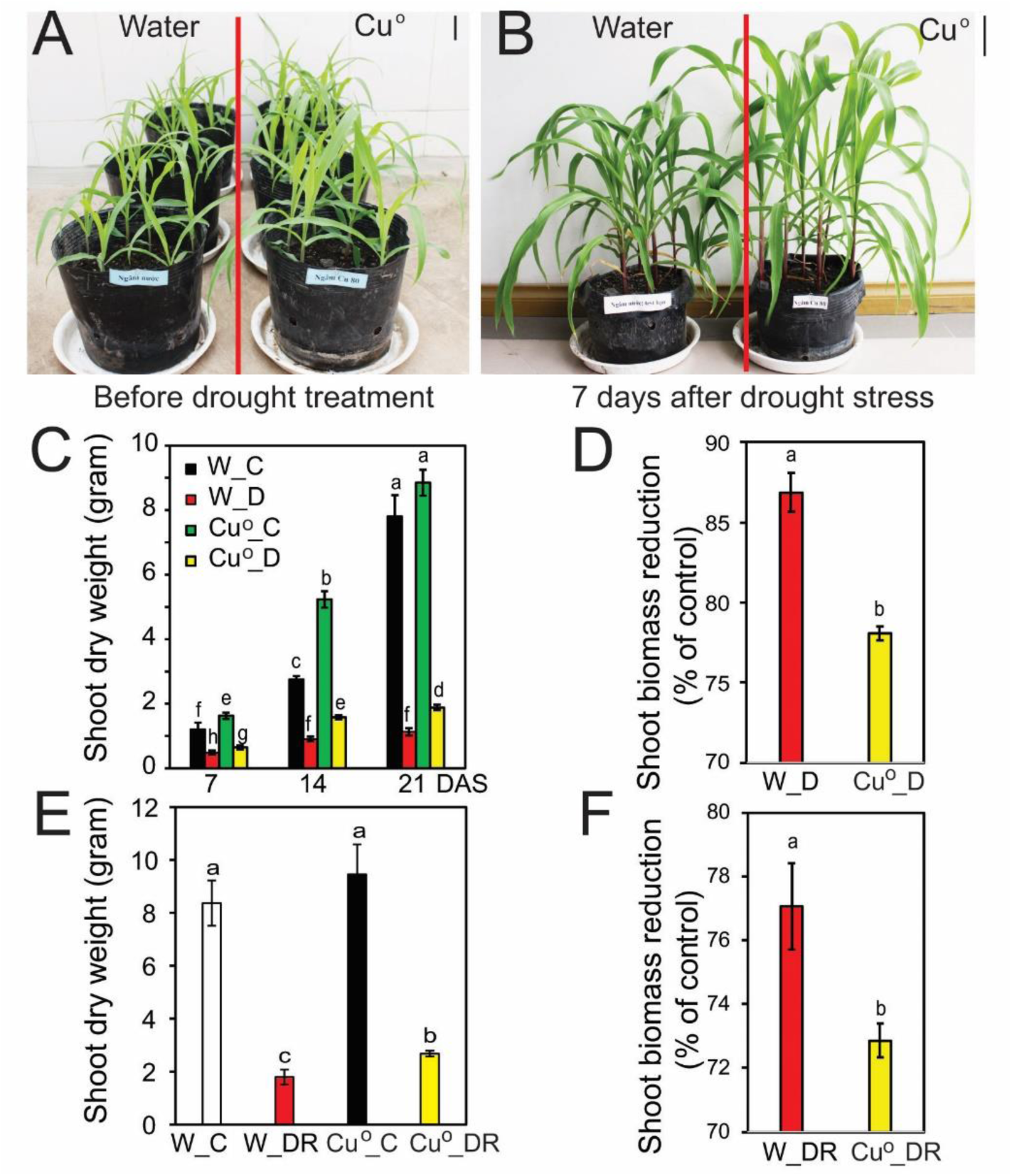
Drought tolerant phenotype of the copper nanoparticle priming in maize. (A) Representative of 12-day-old copper nanoparticle- and water-treated plants were grown under normal conditions. Bar = 2 cm. (B) Representative of copper nanoparticle- and water-treated plants were exposed to drying soil treatment for 7 days. Bar = 3 cm. (C) Shoot dry weight of copper nanoparticle- and water-treated plants at 7, 14 and 21 days after drought stress. (D) Biomass reduction of copper nanoparticle- and water-treated plants at 21 days after drought stress relative to respective well-watered plants. (E) Shoot dry weight of copper nanoparticle- and water-treated plants were exposed to drought for 21 days after drought stress and then re-watered for 7 days. (F) Biomass reduction of copper nanoparticle- and water-treated plants at 7 days after recover relative to respective well-watered plants. Data represent the mean and standard errors (*n* = 5). The different letters (a, b, c, d, e, f, g and h) indicate significant differences between treatments, which was calculated by the multiple Duncan’s test (*P* < 0.05). DAS, days after drought stress; W_C, water-treated control conditions; W_D, water-treated drought conditions; Cu°_C, copper nanoparticle-treated control conditions; Cu°_D, copper nanoparticle-treated drought conditions; W_DR, water-treated drought recovery conditions; Cu°_D, copper nanoparticle-treated drought recovery conditions.

Third, to test how nano-Cu° affects plant drought recovery, the shoot biomass was measured at 7 days of re-watering. As showed in Figure 1E, nano-Cu°-treated plants showed greater shoot biomass than water-treated plants after drought treatment followed by re-watering. Similarly, the shoot biomass reduction was lower in nano-Cu°-treated plants than water-treated plants after drought recovery as compared with non-stressed plants (Figure 1F). Collectively, these results indicated that application of nano-Cu° increased drought tolerance in maize, suggesting that nano-Cu° has a positive effect on drought responses.

### Cu nanoparticle priming maintains maize grain yield under drought

Cu plays an important role in plant growth and development, which can be effect on plant productivity (Karamanos et al. 2004; Xue et al. 2014). We next investigate the roles of nano-Cu° in plant productivity by conducting a yield test assay of maize under non-stressed and drought condition. As showed in Figure 2A, total seed number per plants was similar in both nano-Cu°- and water-treated plants under non-stressed conditions. However, the average seed dried weight per plant of nano-Cu°-treated plants showed higher than that in water-treated plants under non-stressed conditions (Figure 2B). This data revealed the positive effect on plant productivity of nano-Cu° in maize. Under drought conditions, number of seed and average yield per plant of nano-Cu° -treated plants showed significantly higher than water-treated plants (Figure 2). These data collectively indicated that priming with nano-Cu° could rescue the negative effect of drought on maize productivity.

**Figure 2.**
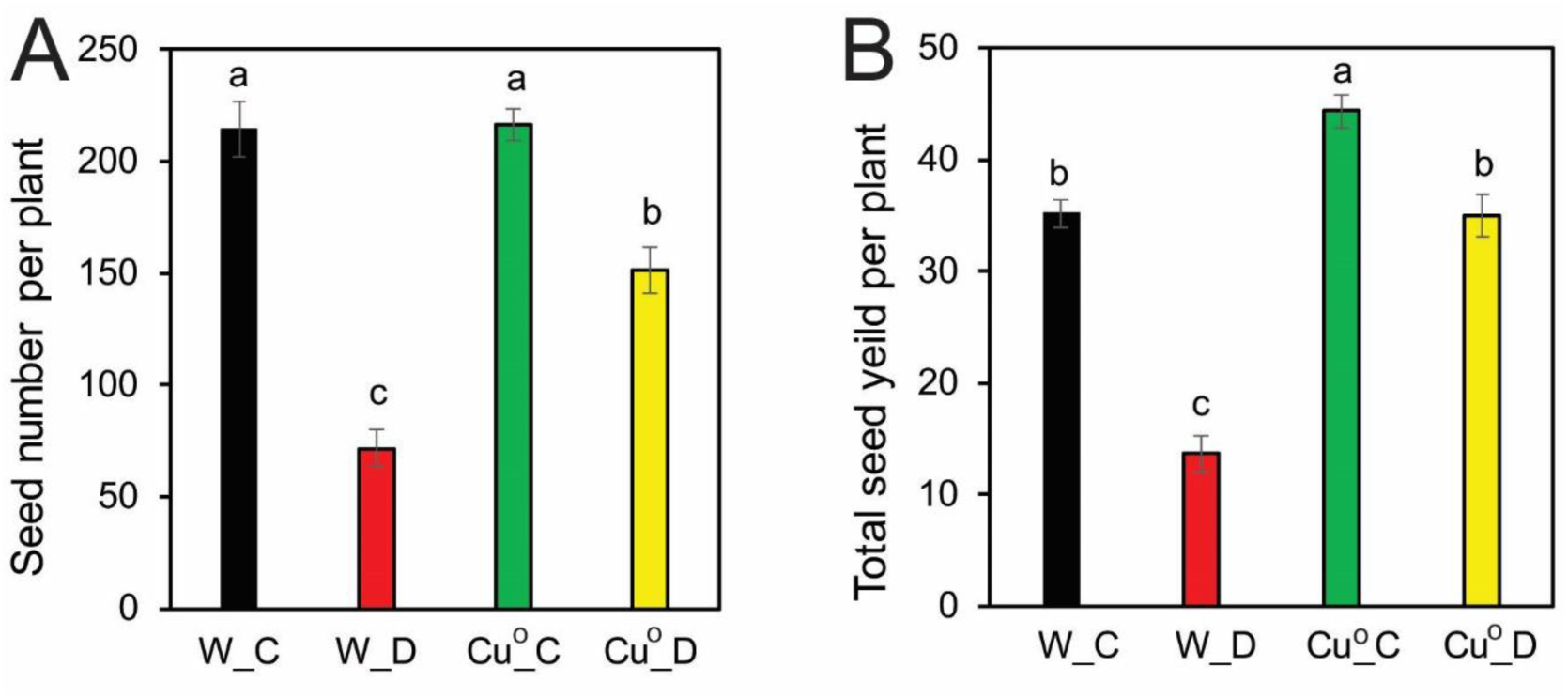
Grain yield of copper nanoparticle-treated and water-treated maize plants under well-watered and drought conditions. Twelve-day-old plants were treated with copper nanoparticle or water for 2 days, then exposed to drought treatment for 21 days. Plants were re-watered and continuously grown under normal condition till the harvesting. (A) Total seed number and (B) seed yield (dried weight) per plant of maize plants treated as described in (A). The seeds were dried for 72 h at 65 °C and measured the dried weight. Data represent the mean and standard errors (*n* = 6). The different letters (a, b and c) indicate significant differences between treatments, which was calculated by the multiple Duncan’s test (*P* < 0.05). W_C, water-treated non-stressed conditions; W_D, water-treated drought conditions; Cu°_C, copper nanoparticle-treated non-stressed conditions; Cu°_D, copper nanoparticle-treated drought conditions.

### Nano-Cu° priming retains relative leaf water status, chlorophyll and carotenoid, and increases anthocyanin contents in maize under drought

We next investigated the effect of nano-Cu° priming on the physiological mechanisms in maize. We found that under non-stress conditions, the levels of RWC of leaf unchanged in nano-Cu°-treated group in comparison with the water-treated group (Figure 3A). Nevertheless, compared with water-treated plant, the nano-Cu°-treated plants revealed higher RWC under drought conditions (Figure 3A). In detail, at 7 days of drought stress, the RWC reduced 7.6 and 11.4 % of nano-Cu°- and water-treated groups, respectively (Figure 3A). At 14 days of drought stress, the RWC significantly declined of 17.3 and 28 % of nano-Cu°- and water-treated plants, respectively (Figure 3A). At 21 days of drought stress, the RWC reduced 30% of nano-Cu°-treated plant and 41.3% of water-treated plants, respectively (Figure 3A). These data indicated that nano-Cu° priming could maintain the water status of maize leaves, therefore reduces negative impact of drought on plant growth, which correlated with greater shoot biomass during drought stress and recovery condition (Figure 1C-F). Additionally, Cu is essential to the growth of plants, which functions in several enzyme processes, and play as a key component in chlorophyll formation (Viera et al. 2019). We next investigate chlorophyll and carotenoid content of maize leaf under nano-Cu° application. Our data showed that there were no significant differences of chlorophyll contents between nano-Cu°-treated and water-treated plants under non-stress conditions (Table 1). In contrast, at 7, 14 and 21 days after drought, respectively, drought caused a sharp decline in the levels of Chl *a* (31, 41 and 61%), Chl *b* (53, 44 and 54%), total Chl (36, 42 and 59%) and carotenoids (19, 36 and 47%) of water-treated plants in drought stress groups, compared with water-treated non-stressed groups (Figure 3B; Table 1). Whereas, nano-Cu°-treated plants in drought stress groups showed reduced the level of Chl *a* (23, 21 and 28%), Chl *b* (26, 29 and 15%), total Chl (23, 23 and 25%) and carotenoids (19, 21 and 36%), compared with nano-Cu°-treated non-stressed plants at 7, 14 and 21 days after drought, respectively (Figure 3B; Table 1). Besides, under drought stress condition, as compared with their respective water-treated plants the levels of Chl *a*, Chl *b*, total Chl, and carotenoids showed higher in nano-Cu°-treated plants than that in water-treated plants (Figure 3B; Table 1). These data indicated that negative effects of drought on the photosynthetic pigments and carotenoids were substantially minimized by exogenous nano-Cu° application. These findings collectively indicated the nano-Cu° priming could mediate recovery on the losses of photosynthetic pigments in drought adaptation in maize (Figure 3B).

**Table 1.**
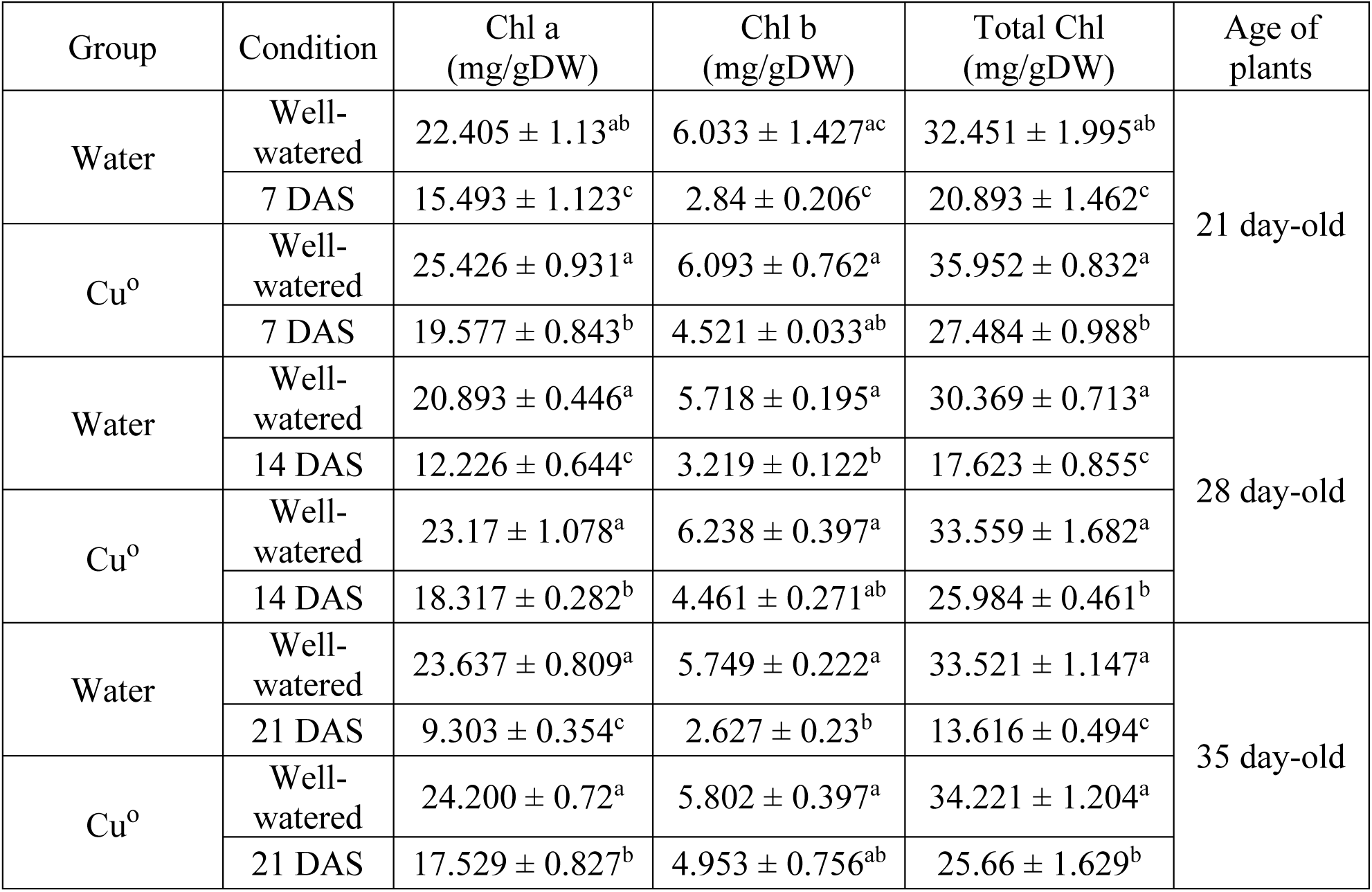
Chlorophyll a, chlorophyll b and total chlorophyll contents of the fifth leaf of copper nanoparticle-treated and water-treated maize plants under well-watered and drought conditions. Data represent the mean and standard errors (*n* = 5). The different letters (a, b and c) indicate significant differences between groups of the same plant age, which was calculated by the multiple Duncan’s test (*P* < 0.05). Chl, chlorophyll; DAS, days after drought stress; DW, dry weight; Cu°, copper nanoparticles.

**Figure 3.**
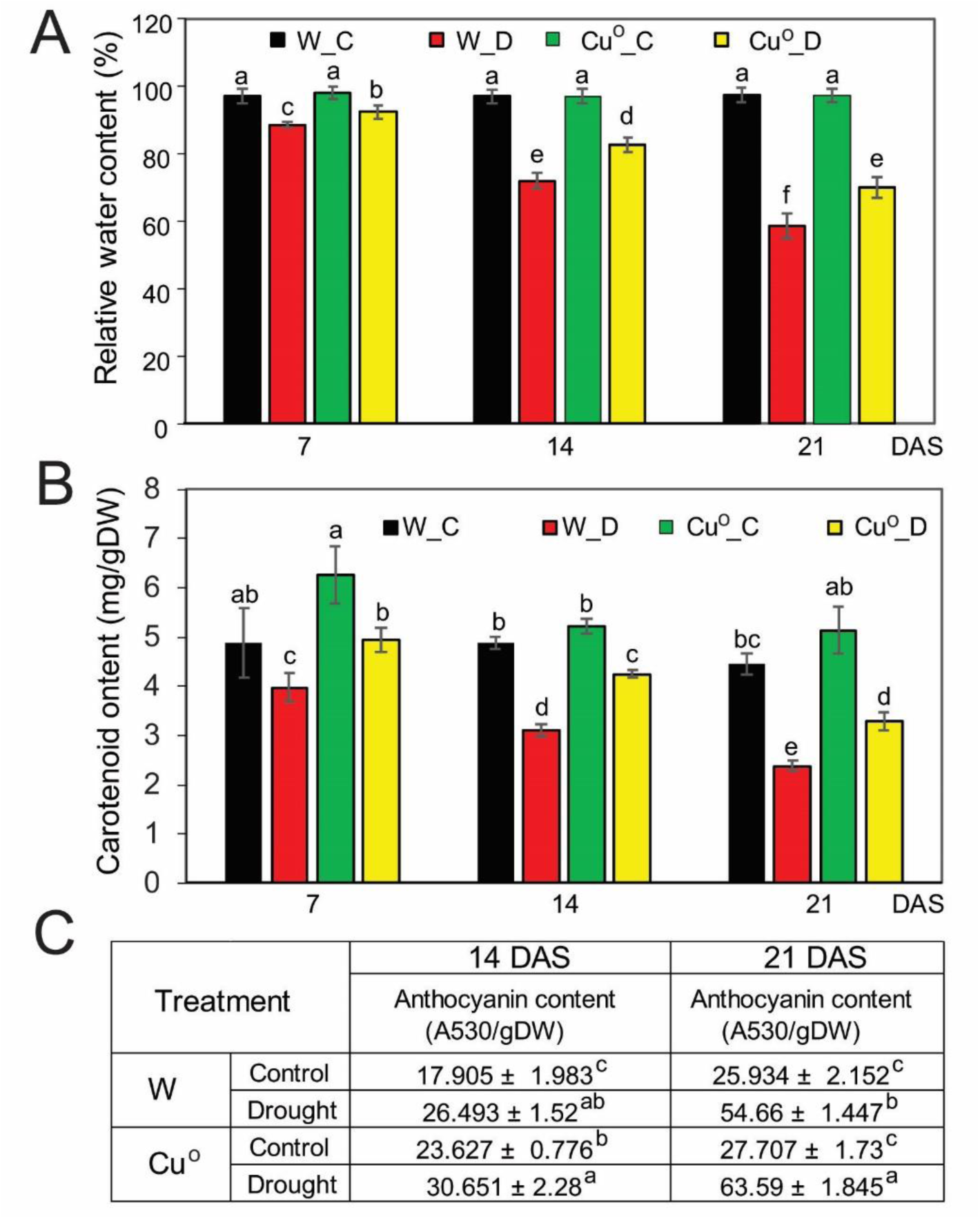
Relative leaf water content, carotenoid and anthocyanin contents of copper nanoparticle-treated and water-treated maize plants under well-watered and drought conditions. (A) Relative water content of the fifth maize leaf under well-watered and drought conditions. Data represent the mean and standard errors (*n* = 5). (B) Carotenoid content of the fifth leaf of maize under well-watered and drought conditions. Data represent the mean and standard errors (*n* = 5). (C) Anthocyanin content of the fifth leaf of maize under well-watered and drought conditions. Data represent the mean and standard errors (*n* = 6). The anthocyanin contents per leaf dry weight were calculated using the ratios of fresh and dry weights of collected leaf parts. The different letters (a, b, c, d, e and f) indicate significant differences between treatments, which was calculated by the multiple Duncan’s test (*P* < 0.05). DAS, days after drought stress; W_C, water-treated non-stressed conditions; W_D, water-treated drought conditions; Cu°_C, copper nanoparticle-treated non-stressed conditions; Cu°_D, copper nanoparticle-treated drought conditions; Cu°, copper nanoparticle.

Several reports have indicated that abiotic stresses, including drought, stimulate anthocyanin production in plants (Li et al. 2017; Nakabayashi et al. 2014; Nguyen et al. 2016). We next investigate the effect of nano-Cu° on anthocyanin accumulation in drought response in maize. We found the anthocyanin content was higher in nano-Cu°-treated plants than water-treated plants (Figure 3C). In detail, anthocyanin content significantly increased in both nano-Cu°- and water-treated plants under drought conditions in comparison with non-stressed plants (Figure 3C). Interestingly, greater anthocyanin content was found in nano-Cu°-treated plants relative to water-treated plants at 21 days of drought stress (Figure 3C). These data indicated that nano-Cu° priming could increase antioxidant contents, which infer an activating of ROS-detoxifying mechanism for drought adaptation in maize.

### Nano-Cu° priming decreases ROS accumulation in maize under drought

In general, to adapt to stresses, plants activate series physiological and biochemical responses that are operated by complex signaling networks (Ha et al. 2014a; Ha et al. 2013a; Mochida et al. 2015; Osakabe et al. 2014; Yamaguchi-Shinozaki and Shinozaki 2006). Under drought stress, plants increase guard cell closure to maintain water status, and enhance activity of antioxidant enzymes to maintain ROS levels in plant cells (Luo et al. 2016; Nguyen et al. 2018b). To determine the potential role of nano-Cu° in ROS homeostasis, we examined the ROS production such as O_2_^•−^ and H_2_O_2_ in the maize leaf of nano-Cu°-treated and control plants under normal and drought stress conditions. We observed that O_2_^•−^ and H_2_O_2_ were accumulated in the leaf of both nano-Cu°-treated and water-treated plants subjected to drought (Figure 4A). However, the accumulation of O_2_^•−^ and H_2_O_2_ nano-Cu°-treated plants were lower levels relative to WT plants (Figure 4A). These results were supported by increased ROS-scavenging enzyme activities such as SOD and APX in nano-Cu° treated plants higher than water-treated under both normal and drought stress conditions, respectively (Figure 4B). In comparison with water-treated plants, activities of SOD and APX were strongly induced in nano-Cu°-treated plants, especially under drought conditions (Figure 4B). In addition, the total soluble protein of leaf of nano-Cu°-treated plants were also higher than that of water-treated plants under drought stress condition (Figure 4B). Collectively, this study indicates that nano-Cu° priming regulates drought response through repression of ROS accumulation in maize plants.

**Figure 4.**
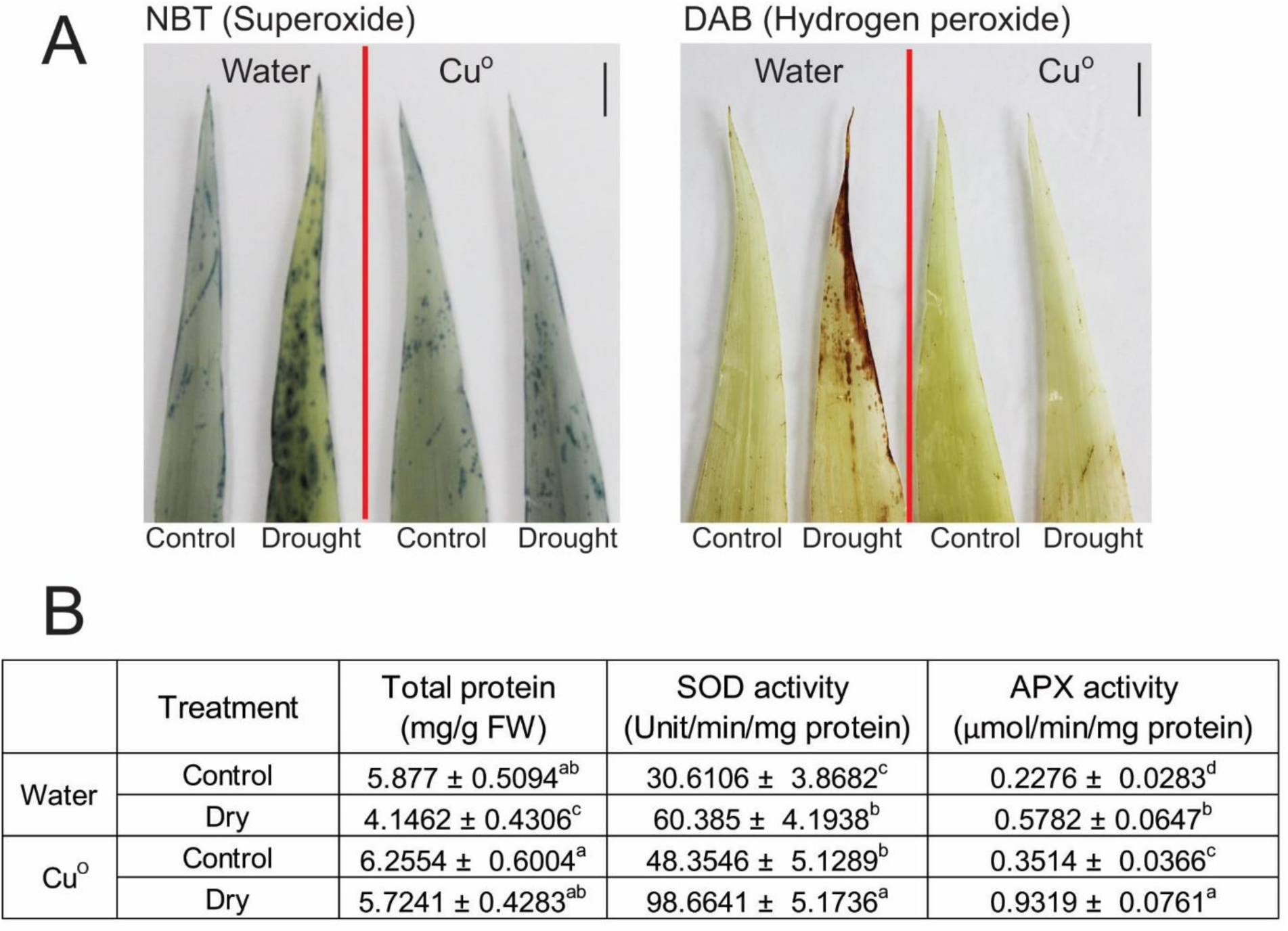
ROS accumulation and ROS-scavenging enzyme activities of copper nanoparticle-treated and water-treated maize plants under well-watered and drought conditions. (A) Histochemical detection of ROS accumulation of the fifth maize leaf under well-watered and at 7 days of drought stress. The nitro blue tetrazolium (NBT) staining detected superoxide and 3,3’-diaminobenzidine (DAB) staining detected hydrogen peroxide. Bar = 1 cm. (B) The total protein, and SOD and APX enzyme activity of the fifth leaf of maize under well-watered and at 7 days after drought stress. Data represent the mean and standard errors (*n* = 5). The different letters (a, b, c and d) indicate significant differences between treatments, which was calculated by the multiple range Duncan’s test (*P* < 0.05). Cu°, copper nanoparticle; FW, fresh weight.

### DISCUSSION

The world population will reach to about 10 billion in 2050, setting a food security for feeding (https://population.un.org/wpp/). However, environmental stresses, particularly drought, negatively affect plant growth and productivity worldwide, caused significant yield losses of crops, including maize (Gong et al. 2014; Hossain et al. 2015; Webber et al. 2018). To cope with environmental stimuli, plants activate multiple adaptive mechanisms from plant growth and development to stress responses (Mochida et al. 2015). Therefore, investigation of agricultural application technology to improve crop productivity is very essential for sustainable human life. In the current study, we provided direct evidences that nano-Cu° priming could enhance drought tolerance in maize, indicated by greater leaf water status (Figure 3A), chlorophyll and carotenoid contents (Figure 3B; Table 1), increased anthocyanin accumulation (Figure 3C) and SOD and APX activities to detoxify the exceed ROS (Figure 4), which may contribute to maintain the photosynthesis and protective mechanism, leading to a growth balancing and drought stress response, could finally contribute to affect plant biomass (Figure 1) and grain productivity (Figure 2) in maize under drought stress condition.

In detail, the positive effect of nano-Cu° priming found in enhancing plant biomass in maize under normal and drought conditions (Figure 1; Supplementary S1). These results were supported by previous studies that Cu plays an important role in plant growth and development, and plant productivity (Karamanos et al. 2004; Ngo et al. 2014; Tamez et al. 2019; Xue et al. 2014). In the present study, the higher plant biomass found in nano-Cu° priming plants indicated the reduction of drought effect on maize, which was associated with the higher water status of leaf in nano-Cu° group (Figure 1C-D and 3A; Supplementary Figure 1). The higher leaf water status of nano-Cu° priming plants could result in maintaining of photosynthesis under drought (Regier et al. 2015; Utsumi et al. 2019), therefore might affect the plant recover and productivity (Figure 1E-F and 2). Support to this hypothesis, our data showed higher chlorophyll contents in nano-Cu° priming plants than those in water-treated plants during drought stress condition (Table 1). Chlorophylls located in chloroplasts have crucial function in photosynthesis system, which highly correlate with plant biomass and recover (Figure 1) and productivity (Figure 2) (Karamanos et al. 2004; Regier et al. 2015; Tamez et al. 2019; Yamauchi 2018). Under drought stress condition, plants accumulate high levels of oxidative stress molecules, including ROS (Farooq et al. 2019), which could directly damage chloroplasts known as the most susceptible organelles to oxidative stress condition (Yamauchi 2018). Our results indicated that the reduction of chlorophyll contents found in both nano-Cu° priming and water-treated groups during drought (Table 1), which may help plants allocating energy and maintain ROS production resulted from photosynthesis to adapt to drought (Casimiro and Arrabaça 2015; Gururani et al. 2015; Huang et al. 2019; Regier et al. 2015; Wang et al. 2018). In plant carotenoid acts as an antioxidant to protect chlorophyll against oxidative stress, therefore maintain chlorophyll content under stress condition (Demmig-Adams and W.Adams III 1996; Emiliani et al. 2018; Latowski et al. 2011). Our results showed that nano-Cu° priming increased carotenoid content during drought in maize (Figure 3B), which could help plant against chlorophyll degradation process, leading to maintain higher chlorophyll contents in nano-Cu° priming plants as compared with water-treated plants (Table 1). Therefore, greater carotenoid content affected by nano-Cu° priming contributed to enhance drought tolerance in maize in this study.

In general, ROS accumulation is increased under stress, which functions in plant defense responses through specific signal transduction pathways (Huang et al. 2019; Xie et al. 2019). Exceed ROS concentrations will increase in cells and cause oxidative damage to membranes, proteins, RNA and DNA molecules, and can even lead to the oxidative destruction of the cell in a process termed oxidative stress (Mittler 2002; Xie et al. 2019). The major sources of ROS during abiotic stress including ROS produced as a consequence of disruptions in metabolic activity and ROS produced for the purpose of signaling as part of the abiotic stress response signal transduction network (Choudhury et al. 2017; Liu and He 2016). Producing high concentration of ROS is harmful to cell, yet they are greatly essential signaling molecules to control stomatal movement at low concentrations to adapt to water deficit (Pei et al. 2000; Wang and Song 2008). In this study, we found that nano-Cu° priming reduced ROS accumulation under drought, which supported to increase drought tolerance in maize via ROS-detoxification mechanism (Figure 4). The steady state of ROS accumulation is crucial for many mechanisms in plant cells, and it is regulated by the balancing between their scavenger activities (Choudhury et al. 2017; Huang et al. 2019). The current study indicated that lower ROS accumulation in nano-Cu° priming plants resulted from increasing activity of SOD and APX enzyme, which helps plants detoxify the exceed ROS, under drought stress condition (Figure 4). These data were supported by previous studies, which reported that applied copper compound nanoparticles increased of the antioxidant systems, including SOD and peroxidase (POD) activities, and total antioxidant levels (Regier et al. 2015; Singh et al. 2017a; Tamez et al. 2019). Signaling ROS directly changes the redox status of multiple regulatory proteins, and alter transcription and translation resulting in the activation of an acclimation response that would mitigate plant stress effects on metabolisms and reduce the level of metabolically produced ROS (Gururani et al. 2015; Liu and He 2016; Xie et al. 2019). In addition, nano-Cu° priming increased anthocyanin content in maize during drought stress (Figure 3C), suggesting that nano-Cu° priming not only affects the ROS-scavenging enzyme activity but also elevates the antioxidant biosynthesis in drought response in maize. Previous studies showed that anthocyanin accumulates during drought, which could help detoxifying exceed ROS for plant stress adaptation (Huang et al. 2019; Li et al. 2017; Nakabayashi et al. 2014). Therefore, greater anthocyanin content affected by nano-Cu° priming contributed to enhance drought tolerance in maize in this study. Consequently, nano-Cu° priming in maize increased both ROS-scavenging enzyme activity and antioxidants to detoxifying exceed ROS molecules in response to drought, leading to a protection of chlorophyll pigments, which could finally maintain grain yield in maize (Figure 2).

## CONCLUSION

Our comparative analyses demonstrated valuable insights into the roles of the nano-Cu° that regulate the plant growth and development as well as drought stress response in maize (Figure 5). Nano-Cu° positively affects plant biomass under both normal and drought condition. Additionally, application of nano-Cu° retains leaf water status, chlorophyll and carotenoid content under drought stress condition. Nano-Cu° priming has a major role in controlling ROS level in maize in response to water stress through increasing both ROS-scavenging enzyme activity and antioxidants. Taken together, nano-Cu° priming has the ability to positively regulate multiple protective mechanisms associated with drought tolerance (Figure 5). Importantly, we provide evidence for maintain maize productivity under drought (Figure 2), which could potentially use for sustainable agriculture application.

**Figure 5.**
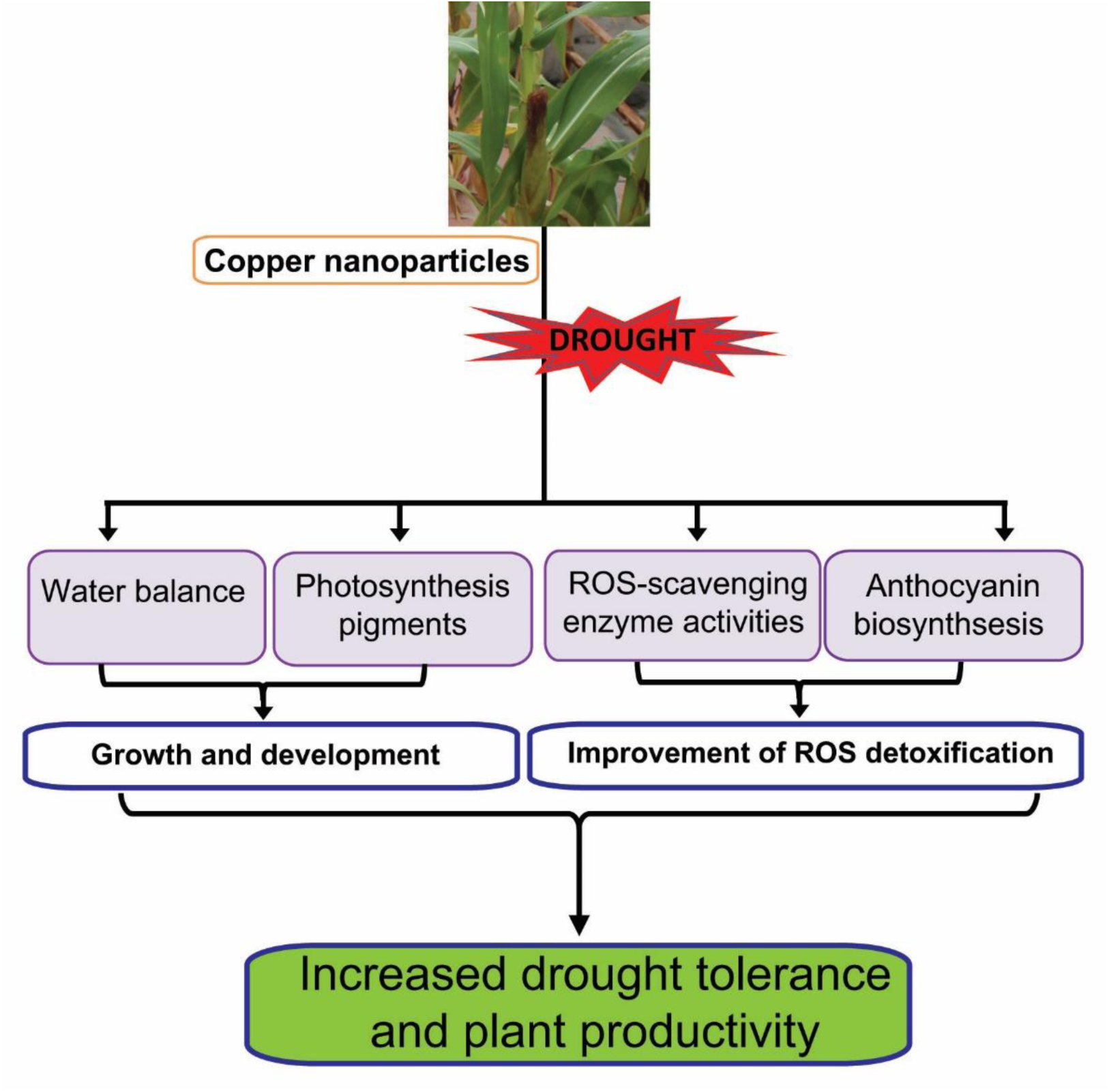
Model for positive regulatory role of copper nanoparticles in response of maize to drought. Copper nanoparticle priming in maize results in changes in various physiological and biochemical processes, including maintained water status of leaves and photosynthesis pigments, and improvement of ROS detoxification through increasing both enzymatic and non-enzymatic antioxidants, which finally increased plant productivity under drought.

## Acknowledgements

This project was supported by grants from project coded KHCN-TB.10C/13-18 to SH under the program titled “Science and Technology for the Sustainable Development of the Northwest Region” of Vietnam National University, Hanoi, Vietnam.

## Author contributions

DN and CH designed research; CH, NL, KN, HL, AN, and ND, performed research; DN and SH contributed research materials, reagents and analytic tools; CH, NL, and HN analyzed the data; and HN, CH and DN wrote the manuscript.

## REFERENCES

Ambrosini VG et al. (2018) High copper content in vineyard soils promotes modifications in photosynthetic parameters and morphological changes in the root system of ‘Red Niagara’ plantlets. Plant Physiol Biochem 128:89–98. https://doi.org/10.1016/j.plaphy.2018.05.011

Arora S, Sharma P, Kumar S, Nayan R, Khanna PK, M.G.H Z (2012) Gold-nanoparticle induced enhancement in growth and seed yield of *Brassica juncea*. Plant Growth Regulation 66:303–310. https://doi.org/10.1007/s10725-011-9649-z

Bradford MM (1976) A rapid and sensitive method for the quantitation of microgram quantities of protein utilizing the principle of protein-dye binding. Anal Biochem 72:248–254. https://doi.org/10.1016/0003-2697(76)90527-3

Burke DJ et al. (2015) Iron oxide and titanium dioxide nanoparticle effects on plant performance and root associated microbes. Int J Mol Sci 16:23630–23650. https://doi.org/10.3390/ijms161023630.

Casimiro A, Arrabaça MC (2015) Effect of copper deficiency on photosynthesis in wheat. In: Sybesma C. (eds) Advances in Photosynthesis Research. Advances in Agricultural Biotechnology, vol 4. Springer, Dordrecht, pp:435–437. https://doi.org/10.1007/978-94-017-4971-8_95

Choudhury FK, Rivero RM, Blumwald E, Mittler R (2017) Reactive oxygen species, abiotic stress and stress combination. Plant J 90:856–867. https://doi.org/10.1111/tpj.13299

Demmig-Adams B, W.Adams III W (1996) The role of xanthophyll cycle carotenoids in the protection of photosynthesis. Trends Plant Sci 1:21–26. https://doi.org/10.1016/S1360-1385(96)80019-7.

Din MI, Arshad F, Hussain Z, Mukhtar M (2017) Green adeptness in the synthesis and stabilization of copper nanoparticles: Catalytic, antibacterial, cytotoxicity, and antioxidant activities Nanoscale Res Lett 12:638. https://doi.org/10.1186/s11671-017-2399-8

Drazkiewicz M, Skorzynska-Polit E, Krupa Z (2004) Copper-induced oxidative stress and antioxidant defence in *Arabidopsis thaliana*. Biometals 17:379–387. https://doi.org/10.1023/B:BIOM.0000029417.18154.22

Emiliani J, D’Andrea L, Lorena Falcone Ferreyra M, Maulion E, Rodriguez E, Rodriguez-Concepcion M, Casati P (2018) A role for beta,beta-xanthophylls in Arabidopsis UV-B photoprotection. J Exp Bot 69:4921–4933. https://doi.org/10.1093/jxb/ery242

Farooq MA et al. (2019) Acquiring control: The evolution of ROS-Induced oxidative stress and redox signaling pathways in plant stress responses. Plant Physiol Biochem 141:353–369. https://doi.org/10.1016/j.plaphy.2019.04.039

Gong F, Yang L, Tai F, Hu X, Wang W (2014) “Omics” of maize stress response for sustainable food production: opportunities and challenges. Omics 18:714–732. https://doi.org/10.1089/omi.2014.0125

Gururani MA, Venkatesh J, Tran LS (2015) Regulation of photosynthesis during abiotic stress-Induced photoinhibition. Mol Plant 8:1304–1320. https://doi.org/10.1016/j.molp.2015.05.005

Ha CV et al. (2014a) Genome-wide identification and expression analysis of the CaNAC family members in chickpea during development, dehydration and ABA treatments. PLoS One 9:e114107 https://doi.org/10.1371/journal.pone.0114107

Ha CV et al. (2013a) The auxin response factor transcription factor family in soybean: genome-wide identification and expression analyses during development and water stress. DNA Res 20:511–524. https://doi.org/10.1093/dnares/dst027

Ha CV, L. DT, Nishiyama R, Watanabe Y, Tran UT, Dong NV, Tran LS (2013b) Characterization of the newly developed soybean cultivar DT2008 in relation to the model variety W82 reveals a new genetic resource for comparative and functional genomics for improved drought tolerance. Biomed Res Int 2013:759657. https://doi.org/10.1155/2013/759657

Ha CV et al. (2014b) Positive regulatory role of strigolactone in plant responses to drought and salt stress Proc Natl Acad Sci U S A 111:851–856. https://doi.org/10.1073/pnas.1322135111

Hossain MA et al. (2015) Hydrogen peroxide priming modulates abiotic oxidative stress tolerance: insights from ROS detoxification and scavenging. Front Plant Sci 6:420 https://doi.org/10.3389/fpls.2015.00420

Huang H, Ullah F, Zhou DX, Yi M, Zhao Y (2019) Mechanisms of ROS regulation of plant development and stress responses. Front Plant Sci 10:800. https://doi.org/10.3389/fpls.2019.00800

Karamanos RE, Pomarenski Q, Goh TB, Flore NA (2004) The effect of foliar copper application on grain yield and quality of wheat. Canadian Journal of Plant Science 84:47–56. https://doi.org/10.4141/P03-090.

Latowski D, Kuczynska P, Strzalka K (2011) Xanthophyll cycle--a mechanism protecting plants against oxidative stress. Redox Rep 16:78–90. https://doi.org/10.1179/174329211x13020951739938

Li P, Li YJ, Zhang FJ, Zhang GZ, Jiang XY, Yu HM, Hou BK (2017) The *Arabidopsis* UDP-glycosyltransferases UGT79B2 and UGT79B3, contribute to cold, salt and drought stress tolerance via modulating anthocyanin accumulation. Plant J 89:85–103. https://doi.org/10.1111/tpj.13324

Liu Y, He C (2016) Regulation of plant reactive oxygen species (ROS) in stress responses: learning from AtRBOHD. Plant Cell Rep 35:995–1007. https://doi.org/10.1007/s00299-016-1950-x

Luo P et al. (2016) Overexpression of Rosa rugosa anthocyanidin reductase enhances tobacco tolerance to abiotic stress through increased ROS scavenging and modulation of ABA signaling. Plant Sci 245:35–49. https://doi.org/10.1016/j.plantsci.2016.01.007

Mittler R (2002) Oxidative stress, antioxidants and stress tolerance. Trends Plant Sci 7:405–410 https://doi.org/10.1016/S1360-1385(02)02312-9

Mochida K, Ha CV, Sulieman S, Nguyen DV, Tran L (2015) Databases of transcription factors in legumes In: de Bruijn FJ (Ed) Biological Nitrogen Fixation, John Wiley & Sons, Inc, New York, pp :817–822. https://doi.org/10.1002/9781119053095.ch81

Mostofa MG, Hossain MA, Fujita M, Tran LS (2015) Physiological and biochemical mechanisms associated with trehalose-induced copper-stress tolerance in rice. Sci Rep 5:11433 https://doi.org/10.1038/srep11433

Myers SS et al. (2017) Climate Change and Global Food Systems: Potential impacts on food security and undernutrition. Annu Rev Public Health 38:259–277. https://doi.org/10.1146/annurev-publhealth-031816-044356

Nakabayashi R et al. (2014) Enhancement of oxidative and drought tolerance in *Arabidopsis* by overaccumulation of antioxidant flavonoids. Plant J 77:367–379. https://doi.org/10.1111/tpj.12388

Ngo BQ, Dao TT, Nguyen CH, Tran XT, Nguyen TV, Khuu TD, Huynh TH (2014) Effects of nanocrystalline powders (Fe, Co and Cu) on the germination, growth, crop yield and product quality of soybean (Vietnamese species DT-51) Advances in Natural Sciences: Nanoscience and Nanotechnology 5:015016. https://iopscience.iop.org/article/10.1088/2043-6262/5/1/015016. Accessed 28 February 2014.

Nguyen HM et al. (2017) Ethanol enhances high-salinity stress tolerance by detoxifying reactive oxygen species in *Arabidopsis thaliana* and rice. Front Plant Sci 8:1001. https://doi.org/10.3389/fpls.2017.01001

Nguyen HM et al. (2018a) Transcriptomic analysis of *Arabidopsis thaliana* plants treated with the Ky-9 and Ky-72 histone deacetylase inhibitors. Plant Signal Behav 13:e1448333. https://doi.org/10.1080/15592324.2018.1448333

Nguyen KH et al. (2016) *Arabidopsis* type B cytokinin response regulators ARR1, ARR10, and ARR12 negatively regulate plant responses to drought. Proc Natl Acad Sci U S A 113:3090–3095 https://doi.org/10.1073/pnas.1600399113

Nguyen KH et al. (2018b) The soybean transcription factor GmNAC085 enhances drought tolerance in *Arabidopsis*. Environmental and Experimental Botany 151:12–20 https://doi.org/10.1016/j.envexpbot.2018.03.017

Osakabe Y, Osakabe K, Shinozaki K, Tran LS (2014) Response of plants to water stress. Front Plant Sci 5:86. https://doi.org/10.3389/fpls.2014.00086

Pei ZM et al. (2000) Calcium channels activated by hydrogen peroxide mediate abscisic acid signalling in guard cells. Nature 406:731–734. https://doi.org/10.1038/35021067

Piperno DR, Ranere AJ, Holst I, Iriarte J, Dickau R (2009) Starch grain and phytolith evidence for early ninth millennium B.P. maize from the Central Balsas River Valley, Mexico. Proc Natl Acad Sci U S A 106:5019–5024. https://doi.org/10.1073/pnas.0812525106

Printz B et al. (2016) Combining -Omics to Unravel the Impact of Copper Nutrition on Alfalfa (*Medicago sativa*) Stem Metabolism. Plant Cell Physiol 57:407–422. https://doi.org/10.1093/pcp/pcw001

Regier N, Cosio C, von Moos N, Slaveykova VI (2015) Effects of copper-oxide nanoparticles, dissolved copper and ultraviolet radiation on copper bioaccumulation, photosynthesis and oxidative stress in the aquatic macrophyte Elodea nuttallii. Chemosphere 128:56–61. https://doi.org/10.1016/j.chemosphere.2014.12.078

Salemi FE, M.N.; Tran, L.S. (2019) Mechanistic insights into enhanced tolerance of early growth of alfalfa (*Medicago sativa* L.) under low water potential by seed-priming with ascorbic acid or polyethylene glycol solution. Industrial Crops and Products 137:436–445 https://doi.org/10.1016/j.indcrop.2019.05.049

Savvides A, Ali S, Tester M, Fotopoulos V (2016) Chemical priming of plants against multiple abiotic stresses: Mission possible?. Trends Plant Sci 21:329–340. https://doi.org/10.1016/j.tplants.2015.11.003

Singh A, Singh NB, Hussain I, Singh H (2017a) Effect of biologically synthesized copper oxide nanoparticles on metabolism and antioxidant activity to the crop plants *Solanum lycopersicum* and *Brassica oleracea* var. botrytis. Journal of Biotechnology 262:11–27 https://doi.org/10.1016/j.jbiotec.2017.09.016

Singh T, Shukla S, Kumar P, Wahla V, Bajpai VK (2017b) Application of nanotechnology in food science: Perception and overview. Front Microbiol 8:1501. https://doi.org/10.3389/fmicb.2017.01501

Stevanovic M et al. (2016) The impact of high-end climate change on agricultural welfare. Sci Adv 2:e1501452. https://doi.org/10.1126/sciadv.1501452

Tamez C, Morelius EW, Hernandez-Viezcas JA, Peralta-Videa JR, Gardea-Torresdey J (2019) Biochemical and physiological effects of copper compounds/nanoparticles on sugarcane (*Saccharum officinarum*). Sci Total Environ 649:554–562. https://doi.org/10.1016/j.scitotenv.2018.08.337

Thangavelu RM, Gunasekaran D, Jesse MI, S.U. MR, Sundarajan D, Krishnan K (2018) Nanobiotechnology approach using plant rooting hormone synthesized silver nanoparticle as “nanobullets” for the dynamic applications in horticulture – an in vitro and ex vitro study. Arabian Journal of Chemistry 11:48–61. https://doi.org/10.1016/j.arabjc.2016.09.022

Utsumi Y et al. (2019) Acetic acid treatment enhances drought avoidance in cassava (*Manihot esculenta* Crantz). Front Plant Sci 10:521. https://doi.org/10.3389/fpls.2019.00521

Viera I, Perez-Galvez A, Roca M (2019) Green natural colorants. Molecules 24 https://doi.org/10.3390/molecules24010154

Wang P, Song CP (2008) Guard-cell signalling for hydrogen peroxide and abscisic acid. New Phyto 178:703–718. https://doi.org/10.1111/j.1469-8137.2008.02431.x

Wang Z et al. (2018) Effects of drought stress on photosynthesis and photosynthetic electron transport chain in young apple tree leaves. Biol Open 7:pii: bio035279. https://doi.org/10.1242/bio.035279

Webber H et al. (2018) Diverging importance of drought stress for maize and winter wheat in Europe. Nat Commun 9:4249. https://doi.org/10.1038/s41467-018-06525-2

Wheeler T, von Braun J (2013) Climate change impacts on global food security. Science 341:508–513. https://doi.org/10.1126/science.1239402

Xie X, He Z, Chen N, Tang Z, Wang Q, Cai Y (2019) The roles of environmental factors in regulation of oxidative stress in plant. Biomed Res Int 2019:9732325. https://doi.org/10.1155/2019/9732325

Xue Y et al. (2014) Zinc, iron, manganese and copper uptake requirement in response to nitrogen supply and the increased grain yield of summer maize. PLoS One 9:e93895 https://doi.org/10.1371/journal.pone.0093895

Yamaguchi-Shinozaki K, Shinozaki K (2006) Transcriptional regulatory networks in cellular responses and tolerance to dehydration and cold stresses. Annu Rev Plant Biol 57:781–803 https://doi.org/10.1146/annurev.arplant.57.032905.105444

Yamauchi Y (2018) Integrated chemical control of abiotic stress tolerance using biostimulants. In: Andjelkovic V (Ed) Plant Abiotic Stress and Responses to Climate Change. IntechOPen, London, pp:133–143. https://doi.org/10.5772/intechopen.74214

