## Supplementary material for "Copper nanoparticle application enhances plant growth and grain yield in maize under drought stress conditions"

**SUPPLEMENTARY MATERIALS**

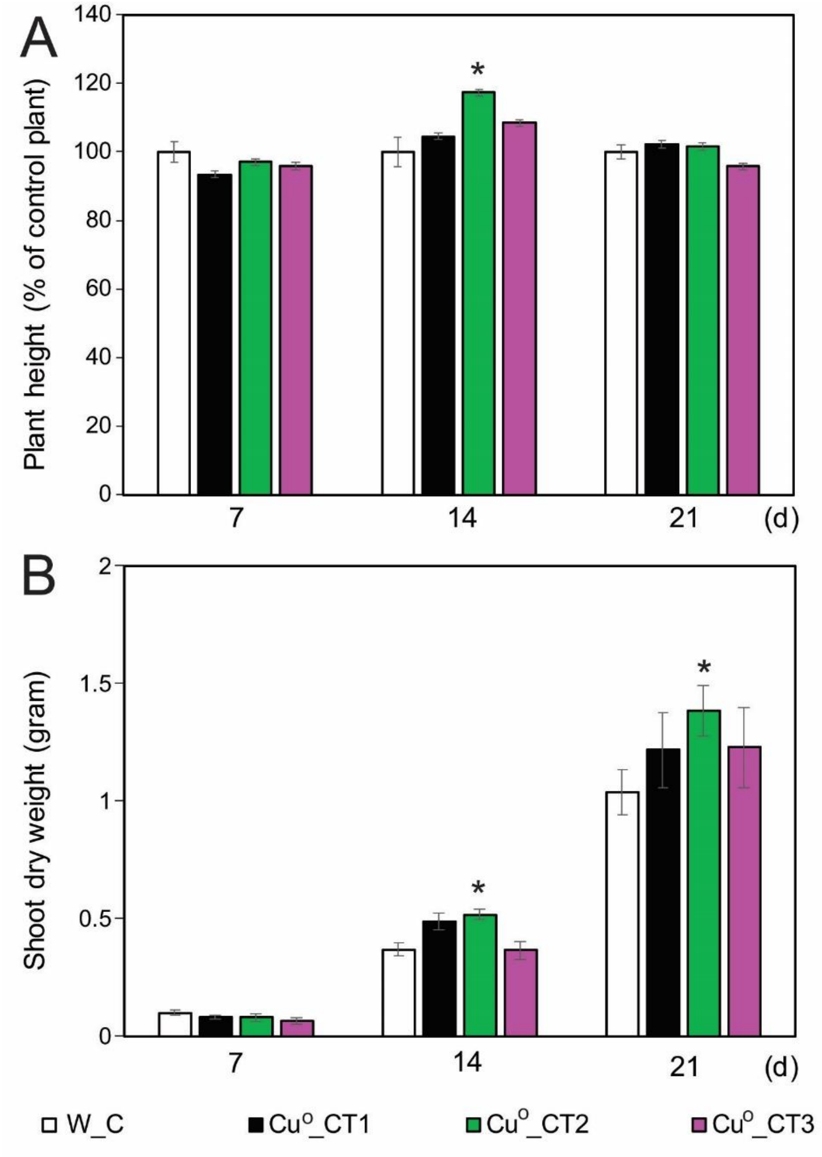


**Supplementary Figure1. Plant height and shoot dry weight of copper nanoparticle-treated and water-treated plants under well-watered growth condition**. (A) Plant height of 7, 14 and 21-day-old copper nanoparticle-treated plants relative to water-treated (control) plants under well-watered condition. Data represent the mean and standard errors ($n$ = 10). Asterisk indicates significant difference between copper nanoparticle-treated and control groups (*$P$ < 0.05). (B) Shoot dry weight of 7, 14 and 21-day-old copper nanoparticle-treated and water-treated (control) plants under well-watered condition. Data represent the mean and standard errors ($n$ = 5). Asterisk indicates significant difference between copper nanoparticle-treated and control groups (*$P$ < 0.05). W_C, plants were treated with water (control); $Cu^o$ _CT1, plants were treated with 52 µM copper nanoparticles; $Cu^o$ _CT2, plants were treated with 69.4 µM copper nanoparticles; $Cu^o$ _CT3, plants were treated with 86.8 µM copper nanoparticles; d, days after sowing seed.
